# Learning to Predict in Networks with Heterogeneous and Dynamic Synapses

**DOI:** 10.1101/2021.05.18.444107

**Authors:** Daniel Burnham, Eric Shea-Brown, Stefan Mihalas

## Abstract

A salient difference between artificial and biological neural networks is the complexity and diversity of individual units in the latter (Tasic et al., 2018). This remarkable diversity is present in the cellular and synaptic dynamics. In this study we focus on the role in learning of one such dynamical mechanism missing from most artificial neural network models, short-term synaptic plasticity (STSP). Biological synapses have dynamics over at least two time scales: a long time scale, which maps well to synaptic changes in artificial neural networks during learning, and the short time scale of STSP, which is typically ignored. Recent studies have shown the utility of such short-term dynamics in a variety of tasks (Masse et al., 2019; Perez-Nieves et al., 2021), and networks trained with such synapses have been shown to better match recorded neuronal activity and animal behavior (Hu et al., 2020). Here, we allow the timescale of STSP in individual neurons to be learned, simultaneously with standard learning of overall synaptic weights. We study learning performance on two predictive tasks, a simple dynamical system and a more complex MNIST pixel sequence. When the number of computational units is similar to the task dimensionality, RNNs with STSP outperform standard RNN and LSTM models. A potential explanation for this improvement is the encoding of activity history in the short-term synaptic dynamics, a biological form of long short-term memory. Beyond a role for synaptic dynamics themselves, we find a reason and a role for their diversity: learned synaptic time constants become heterogeneous across training and contribute to improved prediction performance in feedforward architectures. These results demonstrate how biologically motivated neural dynamics improve performance on the fundamental task of predicting future inputs with limited computational resources, and how learning such predictions drives neural dynamics towards the diversity found in biological brains.

## 1 Introduction

The last decades have witnessed astonishing development in neural network research, both in artificial intelligence (AI) applications to problems across the sciences and in creating model systems for understanding computation in the brain. The essential concepts underlying these advances have their origin in neuroscience: distributed computing using neurons, learning by changing synaptic connections, and hierarchical organization of networks (Fukushima, 1980; Hubel & Wiesel, 1959). Likewise, incorporation of these biological features into AI models, together with analyses of emerging large-scale neural datasets, has allowed these models to inform the underlying biology. However, recent experimental discoveries in neurobiology have outpaced application within computing network models. One of the most salient observations is the heterogeneity of neuronal features showing a remarkably diverse anatomy (Gouwens et al., 2019), gene expression (Tasic et al., 2016), intrinsic dynamics (Teeter et al., 2018), *in vivo* responses (de Vries et al., 2020), and, our focus here, remarkably diverse synaptic properties (Seeman et al., 2018).

In ANNs, updating the connection weights during training represents a mechanism akin to longterm plasticity (Goodfellow et al., 2016). However, neurons also exhibit strong, and strongly diverse, short term dynamics – and these are missing from almost all ANN models. These short-term synaptic plasticity mechanisms (STSP) can be broadly categorized as Short-Term Depression and Short-Term Facilitation. Recent work has shown how STSP can provide networks with short term memory (Masse et al., 2019) and the ability to easily solve detection of change tasks (Hu et al., 2020). In addition, recent work in ANNs has incorporated maintenance of temporary state information by implementing “fast weights” in a standard RNN layer architecture (Ba et al., 2016). The authors accomplish this by constructing a “fast” weight matrix in parallel with the standard weight matrix. The fast weight matrix depends on the correlation of current and past hidden states, and its output contributes additively to the evolution of the network hidden state at each time step. Our approach differs in several interesting ways. Chief among them is that the (fast) synaptic dynamics we study represent a multiplicative adaptation at the level of all synapses emerging from an active neuron, rather than a synapse-specific facilitation of synaptic strength for *pairs* of co-active synapses, as in Ba et al. (2016). A second difference is the method of training the parameters for the short term plasticity, which in our model is based on the same optimization as the synaptic weights.

Here, we build on these results in three major ways. First, we allow the timescale of STSP in each individual neuron to be learned, simultaneously with standard learning of overall synaptic weights, via unified gradient-based tools to minimize task training loss in Pytorch (extending from Hu et al. (2020)). Second, we draw explicit comparison to standard RNN and LSTM models with related, but distinct, mechanisms. And third, we study the role of STSP in prediction, a fundamental temporal computation. Prediction tasks are inspired by predictive coding, which has a long tradition in signal processing (Elias, 1955) and has been studied for more than two decades in computational neuroscience (Rao & Ballard, 1999; Huang & Rao, 2011) as well as more recent work on prediction in the machine learning and ANN literature (e.g. with recurrent convolutional LTSMs (Lotter et al., 2016) and video prediction). While predictive coding training functions have been shown to reproduce several features observed in neuronal data (Rao & Ballard, 1999), there are also differences (Zhuang et al., 2021). Here, we explore the role of STSP in a very simplified dynamical system prediction task as well as a more complex sequential MNIST prediction task.

Very recent work has reached similar scientific conclusions to ours, namely that heterogeneous intrinsic dynamics help temporal integration (Perez-Nieves et al., 2021) but there are key differences which make these studies complementary. First, we focus on STSP dynamics, as opposed to cellular and synaptic integration time constants. Second, we focus on temporal prediction rather than classification. Finally, we work with rate-based networks, which enables us to make comparison with standard RNN and LSTM machine learning tools.

## 2 Results

### 2.1 Task description

We investigated the role of STSP in learning by training a set of neural network models with varying usages of STSP dynamics to predict future sequences in two different task environments: time series prediction of N-dimensional sinusoidal (Figure 1B) and N-dimensional decaying exponential (Appendix 5) dynamics, and prediction of pixel intensity in sequentially read MNIST images (Figure 3A).

**Figure 1:**
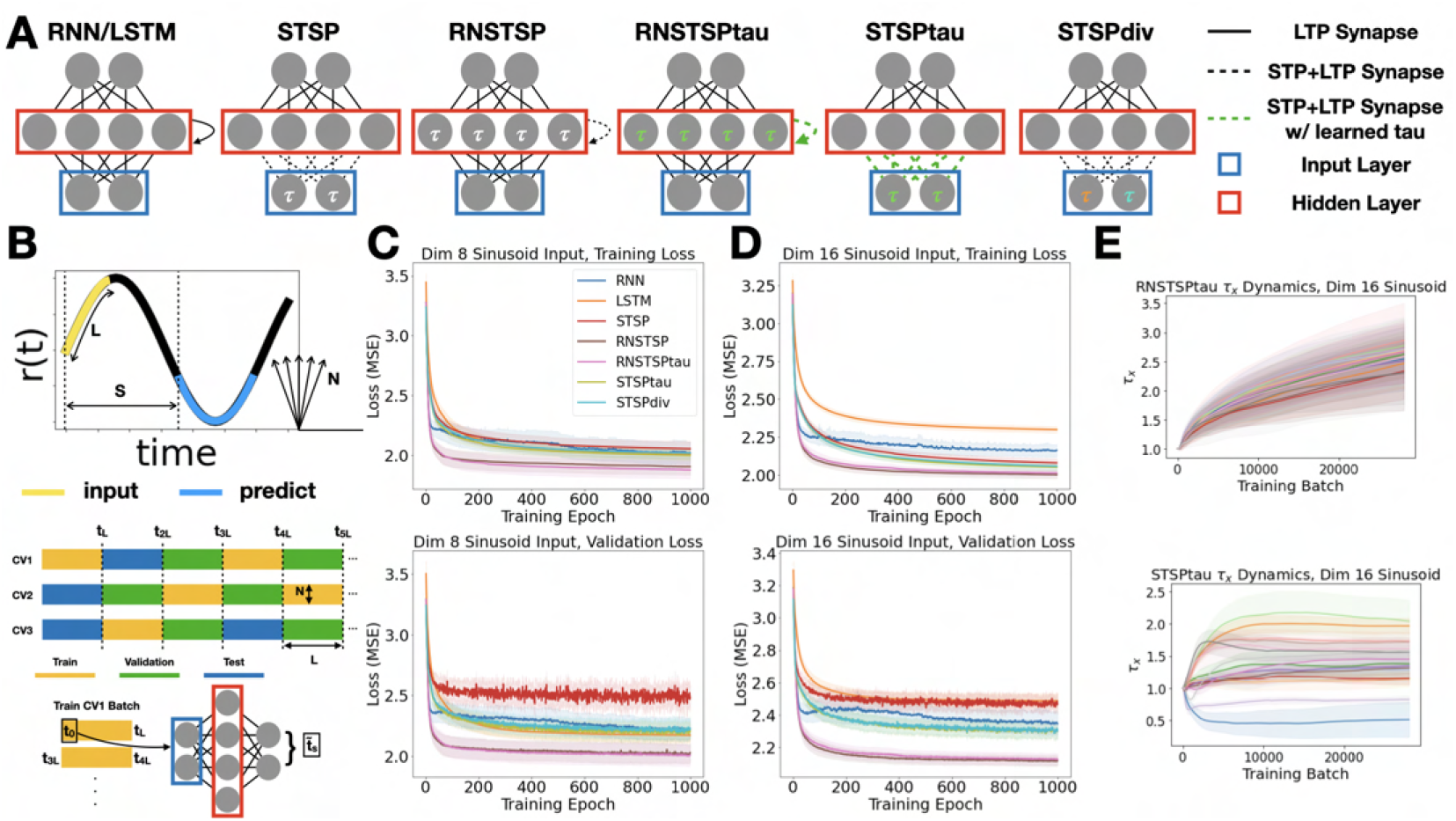
**A** The network models. White parameters are static and homogeneous. Green parameters denote time constants that are learned. Multicolored parameters are static but heterogeneous. STP denotes short-term plasticity synapses and LTP indicates long-term plasticity connections, as in standard ANN learning. **B** Models were trained to perform a time series prediction task on an N- dimensional dynamic. The number of steps ahead to predict is S and the input sequence length is L. Time series train/validation/test split strategy randomly assigns sections of length L to each set. **C** Training and validation loss for parameter matched models predicting an 8 dimensional sinusoid. **D** Training and validation loss for parameter matched models predicting an 16 dimensional sinusoid. **E** Dynamics of STSP time constants for the RNSTSPtau (Top) and STSPtau (Bottom) models predicting a 16 dimension input. Individual lines are time constant values for each STSP node across training.

### 2.2 Model description

We implement a model for short-term depression (Dayan & Abbott, 2001), similar to Hu et al. (2020), in firing-rate network models. The rate of change of synaptic resources for neuron *i* with short-term depression synapses is given by Equation 1, where *x*_*i*_(*t*) represents the synaptic resources, *U* is a constant, *r*_*i*_(*t*) represents the presynaptic activity at time *t*, and 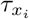 is the time constant of the synapses formed by neuron *i*. The firing rate of a given postsynaptic neuron *j* is represented by Equation 2. Here, *W*_*ij*_ represents the overall synaptic strengths subject to long-term plasticity as in standard ANN learning, and subject to additional modulation on faster timescales via the STSP rule (Hu et al., 2020). The non-linearity, Φ, is a ReLU activation function.

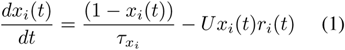

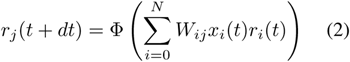

All models have 3-layer architectures with a hidden layer of varying size to facilitate parameter matching across models (Figure 1A). Parameter matching was done by fixing the hidden layer dimension at 16 for the RNN and matching the other models to the corresponding trainable parameter count by modulating the hidden layer dimensions (Section A.2). Short-term synaptic dynamics were included in feedforward models (STSP, STSPtau, STSPdiv) connecting the input layer to hidden layer, as well as in RNNs (RNSTSP, RNSTSPtau) with the STSP synapses within the hidden layer (Figure 1A). These models with short-term plasticity are differentiated in both the initialization of the short-term dynamic time constants and the dynamics of these time constants across training. In the STSP and RNSTSP models, the time constants for each hidden node are homogeneous (*τ*_*x*_ = 3) and static. The STSPtau and RNSTSPtau models likewise have all time constants initialized homogeneously (*τ*_*x*_ = 1), but the *time constants are learned model parameters* trained via the gradient of the task loss, thus permitting the values to be optimized across training. The time constants for the STSPdiv model, by comparison, are static parameters but are initialized at heterogeneous values based on the final learned time constants for the STSPtau model

### 2.3 STSP improves learning of temporal dynamics

The results presented here use sinusoidal input stimuli of dimensions 8 and 16 that have frequencies along each dimension which are evenly spaced across a range of frequencies from 0.001 to 0.333 Hz. Our results indicate that a significant performance advantage is conferred to the RNNs with STSP. This benefit is evidenced by lower minimum loss values for the RNSTSP and RNSTSPtau models across the training and validation sets (Figure 1C, D). A Kruskal-Wallis test shows significant (*P* < .001) differences in the final 200 epochs of the validation loss between the standard RNN and LSTM models and the RNSTSP, and RNSTSPtau models. A post hoc Mann-Whitney test shows significant differences in the validation loss across the final 200 epochs, for both dimension 8 and 16 sinusoidal inputs, between the following model pairs: RNSTSPtau and RNN (*P* < .001), RNSTSP and RNN (*P* < .001), RNSTSPtau and LSTM (*P* < .001), and RNSTSP and LSTM (*P* < .001).

Furthermore, training the time constants in the RNSTSPtau and STSPtau models results in observable diversification (Figure 1E). In addition, possessing a diversified, or diversifiable, set of STSP dynamics differentiates model performance in feedforward model architecture. This differentiation is evidenced by the STSPtau and STSPdiv models significantly outperforming the STSP model (Figure 1C, D). A Kruskal-Wallis test shows significant (*P* < .001) differences in the final 200 epochs of the validation loss between the STSP, STSPtau, and STSPdiv models. A post hoc Mann-Whitney test shows significant differences between the model pairs of STSPtau and STSP (*P* < .001), and STSPdiv and STSP (*P* < .001) for dimension 8 and 16 sinusoidal inputs.

Lastly, when expanding the dimension of the hidden layers of each model to 256 nodes, after 1000 training epochs, the LSTM model achieves the lowest validation loss of all the models for the 4 and 8 dimensional sinusoidal prediction tasks, but does not significantly outperform the RNSTSP and RNSTSPtau models for the 16 dimension sinusoidal prediction (Figure 2). Overfitting is observed for all models with 256 hidden nodes as evidenced by validation loss increasing across training (Figure 2). The RNSTSP and RNSTSPtau models achieve overall minimum validation loss values comparable to the LSTM model. The RNSTSP and RNSTSPtau models achieve this minimum validation loss earlier than the LSTM model in the case of predicting the 4 dimensional sinusoid. When the number of units is much larger than task dimensionality, the RNSTSP and RNSTSPtau models appear especially prone to overfitting. Therefore adopting an early stop criteria during training is necessary to allow these models to continue to perform well in large network implementations.

**Figure 2:**
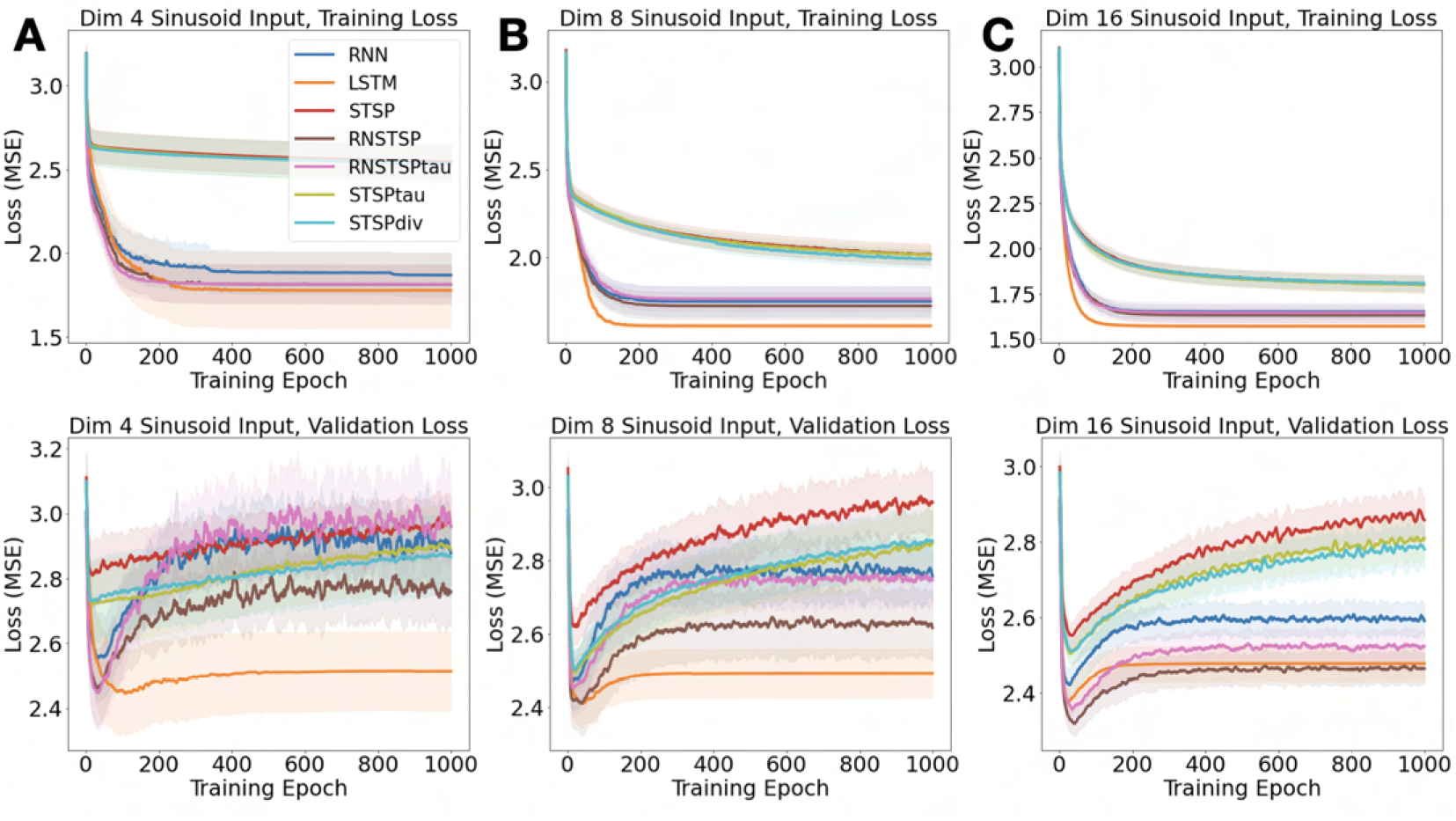
Training and validation loss for models, each with 256 hidden layer nodes, predicting (**A**) 4, (**B**) 8, and (**C**) 16 dimensional sinusoids. Unlike Figure 1, models here are matched for the number of units, resulting in more parameters for LSTM (Table 5). The frequencies along each dimension were again selected to be evenly spaced across a range of frequencies from 0.001 to 0.333 Hz.

### 2.4 STSP improves learning of MNIST pixel sequences

We also compared the performance of our models on a sequential MNIST prediction task. The models were trained on a subset of the full LeCun & Cortes (2010) MNIST dataset containing only the digits 2 and 8 to create a simple but not trivial task. The images were preprocessed such that the pixel row sequences were stacked along 4 input dimensions. The models were trained to predict 4 steps ahead along the pixel sequence.

Our results again indicate that a significant performance advantage is conferred to the RNNs with STSP (Figure 3B, C). A Kruskal-Wallis test shows significant (*P* < .01) differences in the final 200 epochs of the validation loss between the standard LSTM model and the RNSTSP, and RNSTSPtau models. A post hoc Mann-Whitney test shows significant differences in the final 200 epoch validation loss between the RNSTSPtau and LSTM (*P* < .001), RNSTSP and LSTM (*P* = .008), and RNSTSPtau and RNN (*P* = .053) model pairs. No statistically significant performance differences are observed due to heterogeneity of STSP dynamics.

**Figure 3:**
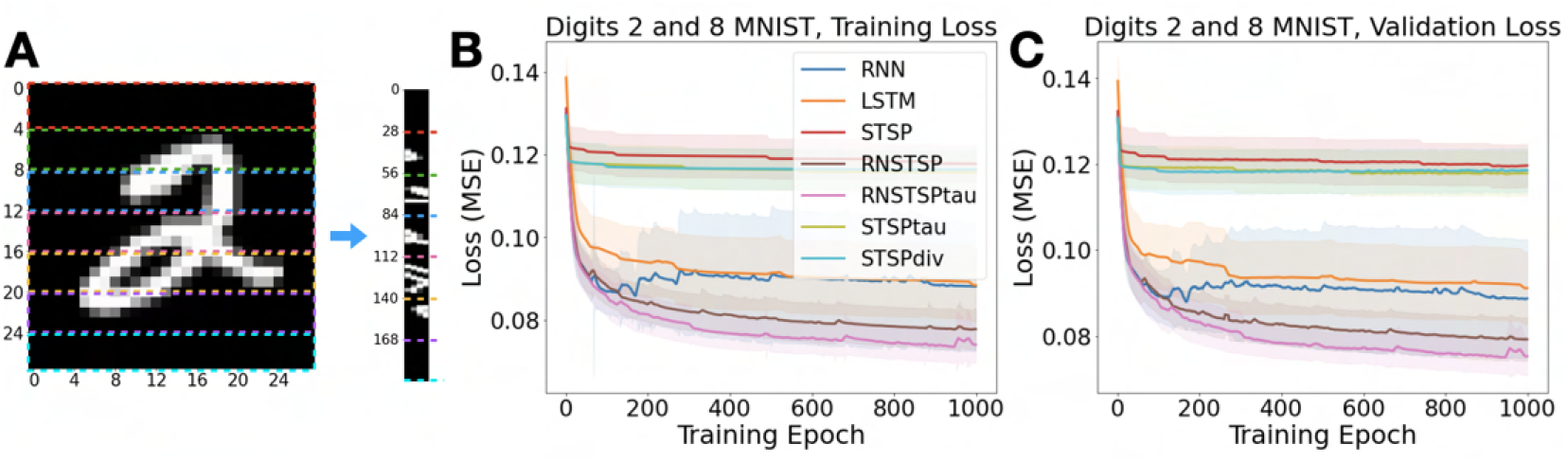
**A** An example transformation of an MNIST image into an MNIST pixel sequence. Four row sections of the original image are horizontally stacked to generate a 4 dimensional sequence. **B** Training and **C** validation loss for predicting 4 steps ahead along the MNIST sequence.

## 3 Discussion

Our results in the first tasks studied, that of predicting a simple dynamical system, demonstrate a performance advantage for RNSTSP (RNNs with short-term synaptic dynamics) over standard RNNs and LSTMs. This holds across a set of predicted dimensions (Figure 1C, D) and dynamics (Section A.3, Figure 5) when the number of computational units is of the same order of magnitude as the task dimensionality. These results are confirmed in the second, more complex task, that of predicting sequential MNIST pixels (Figure 3). When the number of computational units significantly exceeds the task dimensionality, the RNSTSP models learn rapidly (Figure 2) but quickly start to overfit. A potential explanation for RNSTSP performance is the availability of the activity history in the short-term synaptic efficacy. We can view the short-term plasticity as a biological way to implement long short-term memory.

Our second finding is that heterogeneity in the time constants of STSP dynamics, though emergent in both recurrent and feedforward models with learned time constants (Figure 1E), significantly improves learning only in the feedforward setting (Figure 1C D). Recurrent architectures may already allow nodes to manufacture different time scales of activity through the arbitrarily long or short activity trajectories within the recurrent layer. This property could make “ready-made” synaptic timescales, via STSP, a less significant resource for predicting time varying signals. In the feedforward setting, while diversity in these time constants was important, we did not find evidence that they needed to be learned on a neuron-specific basis in tandem with connection weights. Specifically, we found identical performance for the STSPtau model, in which this tandem learning did occur, and the STSPdiv variant, in which the heterogeneous time constants were pre-assigned and held fixed during learning (Figure 1C, D and Figure 3C). This indicates that effective learning of these time constants in nature could be in response to developmental or learning pressures on larger time scales than individual stimulus and task learning. A related idea is that once time constants have been learned to solve a given task, they will likely be of service in solving other tasks with relevant information over similar time scales. This said, the situations in which heterogeneity of synaptic dynamics impacts learning remain to be thoroughly explored. We believe that our findings open doors to future work along these lines, which should analyze temporal tasks which are gradually more complex and more ethologically relevant. To complement this, future efforts should also be made to demonstrate the theoretical underpinnings of how synapses acting as diverse dynamical components can facilitate the representation – and, more importantly, prediction – of time varying inputs.

## Acknowledgments

We wish to thank the Allen Institute founder Paul G Allen for his vision, encouragement and support.

## A Appendix

### A.1 Model Implementation

For both tasks the models were trained using the ADAM optimizer with the default settings (learning rate of 0.001 and beta values of 0.9, 0.999). Models were trained for 1000 epochs with 20 crossvalidation folds using mean squared error loss. Our Pytorch model implementations will be made available to the community through Github.

Of note, for presentation purposes the validation loss plots in Figure 1C, D are smoothed using a moving average with a window size of 3 epochs. The loss trajectories are not qualitatively different without smoothing (Figure 10). Additionally, due to exploding gradients, only 18 CV folds were used for the RNN model predicting the dimension 16 sinusoid in Figure 1C, and 1D due to exploding gradients. Similarly, in Figure 3B, and 3C, only 15, 19, and 19 CV folds were used for the RNN, RNSTSP, and RNSTSPtau models respectively.

### A.2 Model Trainable Parameters

Tables 1-3 detail the hidden layer dimensions and trainable parameter counts for the implemented models. Table 4 details the trainable parameter counts for simulations described in Supplementary Figure 6, for which the number of hidden units was conserved across model type. Table 5 details the trainable parameter counts for simulations described in Figure 2, for which the number of hidden units was set at 256.

**Table 1:** The parameter matched hidden layer sizes and corresponding trainable parameter counts for an input layer dimension of 4. Relevant for models presented in Suplementary Figure 4.

| Model | Hidden Layer Size | Trainable Parameters |
| --- | --- | --- |
| RNN | 16 | 420 |
| LSTM | 7 | 396 |
| STSP | 32 | 301 |
| RNSTSP | 16 | 420 |
| RNSTSPtau | 16 | 436 |
| STSPtau | 32 | 305 |
| STSPdiv | 32 | 301 |

**Table 2:** The parameter matched hidden layer sizes and corresponding trainable parameter counts for an input layer dimension of 8. Relevant for models presented in Figure 1.

| Model | Hidden Layer Size | Trainable Parameters |
| --- | --- | --- |
| RNN | 16 | 552 |
| LSTM | 7 | 540 |
| STSP | 32 | 552 |
| RNSTSP | 16 | 552 |
| RNSTSPtau | 16 | 568 |
| STSPtau | 32 | 569 |
| STSPdiv | 32 | 552 |

**Table 3:** The parameter matched hidden layer sizes and corresponding trainable parameter counts for an input layer dimension of 16. Relevant for models presented in Figure 1.

| Model | Hidden Layer Size | Trainable Parameters |
| --- | --- | --- |
| RNN | 16 | 816 |
| LSTM | 7 | 828 |
| STSP | 24 | 808 |
| RNSTSP | 16 | 816 |
| RNSTSPτ | 16 | 832 |
| STSPτ | 24 | 824 |
| STSPdiv | 24 | 824 |

**Table 4:** Trainable parameter counts for models with matched hidden layers dimensions of 16 nodes.

| Model | Input Dimension | Trainable Parameters |
| --- | --- | --- |
| RNN | 4 | 420 |
| LSTM | 4 | 1476 |
| STSP | 4 | 148 |
| RNSTSP | 4 | 420 |
| RNSTSPτ | 4 | 436 |
| STSPτ | 4 | 152 |
| STSPdiv | 4 | 148 |
| RNN | 8 | 552 |
| LSTM | 8 | 1800 |
| STSP | 8 | 280 |
| RNSTSP | 8 | 552 |
| RNSTSPτ | 8 | 568 |
| STSPτ | 8 | 288 |
| STSPdiv | 8 | 280 |
| RNN | 16 | 816 |
| LSTM | 16 | 2448 |
| STSP | 16 | 544 |
| RNSTSP | 16 | 816 |
| RNSTSPτ | 16 | 832 |
| STSPτ | 16 | 560 |
| STSPdiv | 16 | 544 |

**Table 5:** Trainable parameter counts for models with 256 node hidden layers. Relevant for models presented in Figure 2

| Model | Input Dimension | Trainable Parameters |
| --- | --- | --- |
| RNN | 4 | 68100 |
| LSTM | 4 | 269316 |
| STSP | 4 | 2308 |
| RNSTSP | 4 | 68100 |
| RNSTSPτ | 4 | 68356 |
| STSPτ | 4 | 2312 |
| STSP$\text{div}$ | 4 | 2308 |
| RNN | 8 | 70152 |
| LSTM | 8 | 274440 |
| STSP | 8 | 4360 |
| RNSTSP | 8 | 70152 |
| RNSTSPτ | 8 | 70408 |
| STSPτ | 8 | 4368 |
| STSP$\text{div}$ | 8 | 4360 |
| RNN | 16 | 74256 |
| LSTM | 16 | 284688 |
| STSP | 16 | 8464 |
| RNSTSP | 16 | 74256 |
| RNSTSPτ | 16 | 74512 |
| STSPτ | 16 | 8480 |
| STSP$\text{div}$ | 16 | 8464 |

### A.3 Supplemental Figures

**Figure 4:**
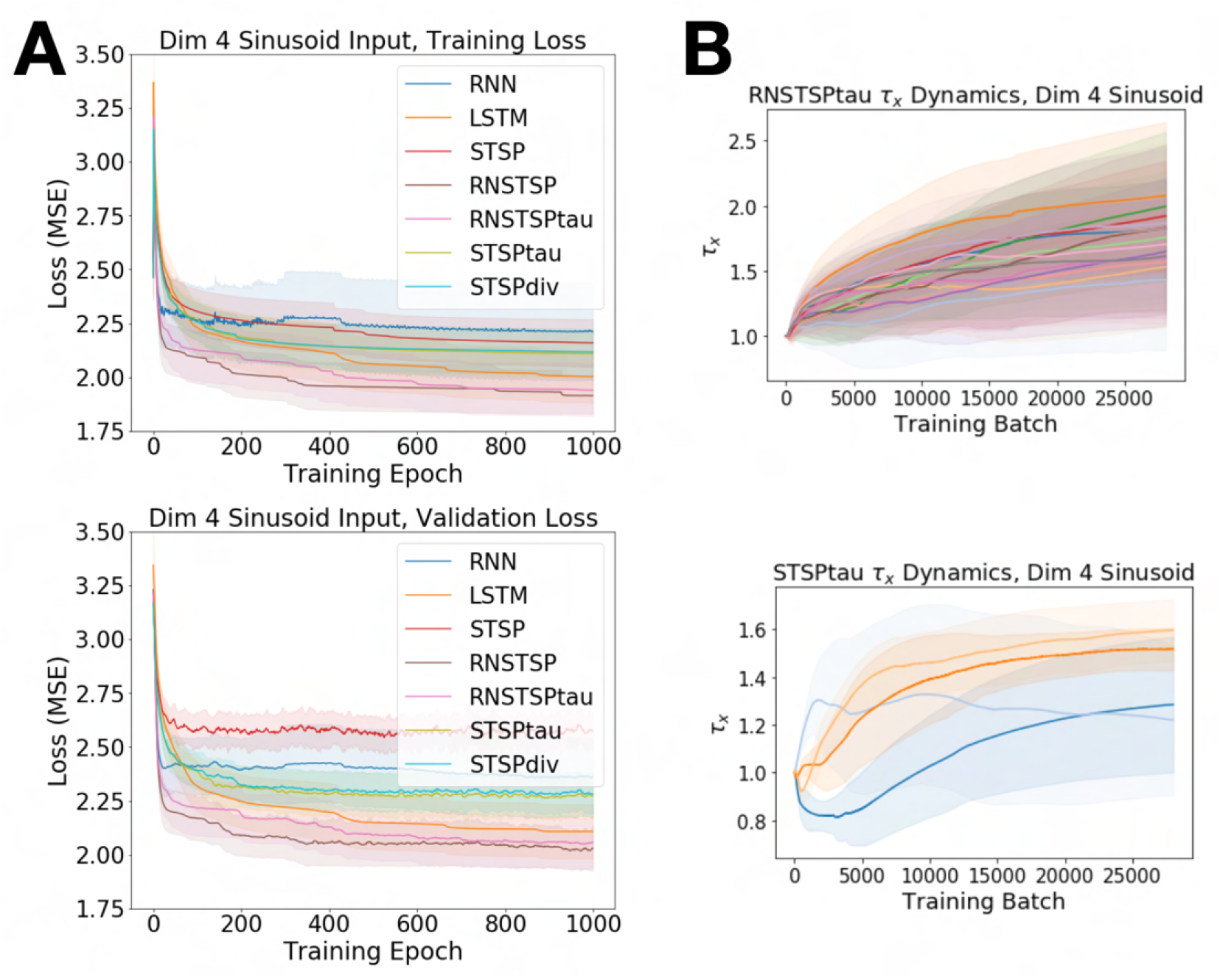
Supplement to Figure 1 demonstrating model performance in a 4 dimensional sinusoid prediction task. **A** The number of hidden units is varied to parameter match the models. Training (Top) and validation (Bottom) loss across training epoch. **B** Heterogeneity of synaptic dynamic time constants for the RNSTSPtau (Top) and STSPtau (Bottom) parameter matched models.

**Figure 5:**
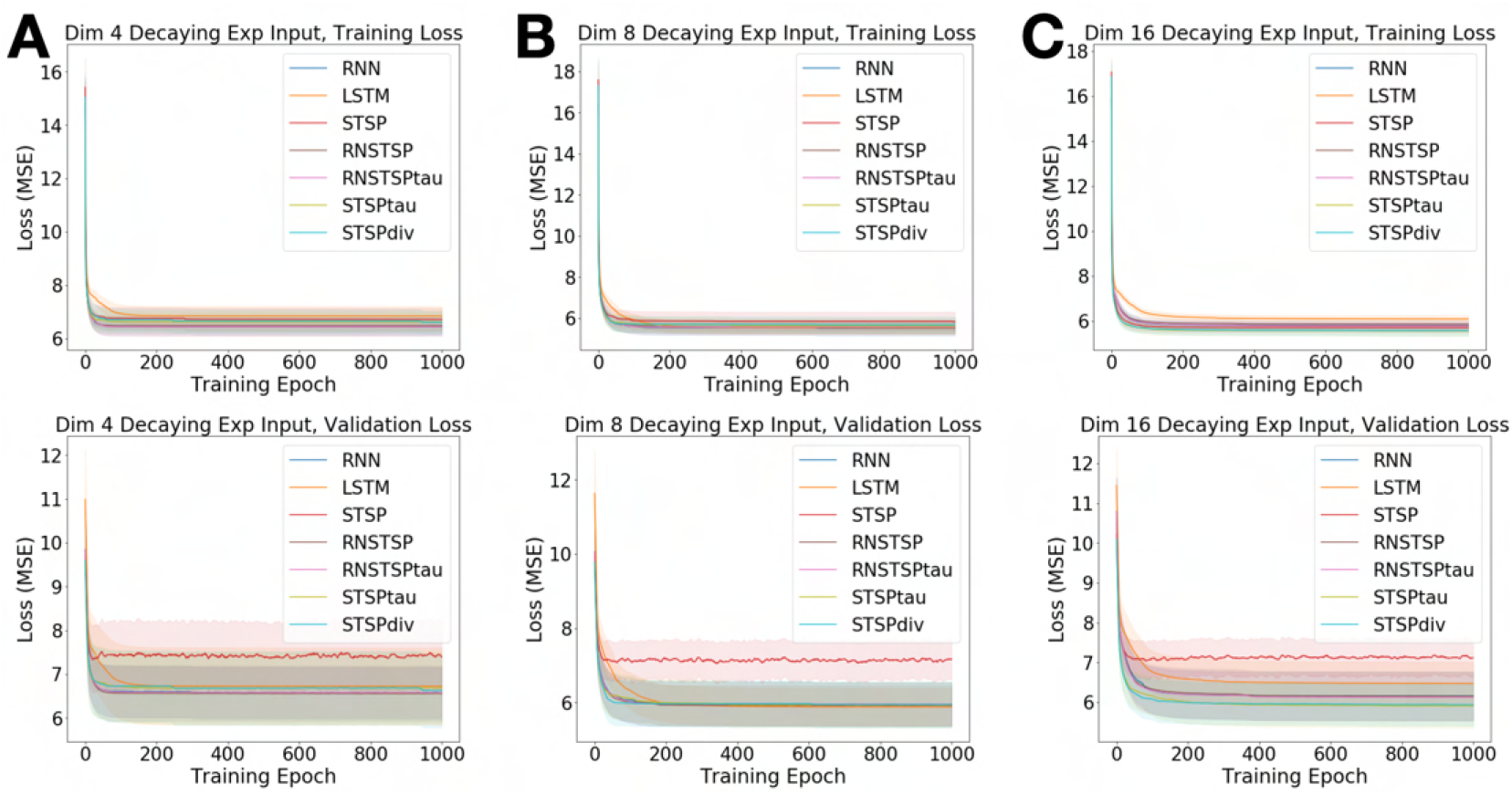
Training (Top) and validation (Bottom) loss across training epoch for models predicting a 4 **A**, 8 **B**, and 16 **C** dimensional decaying exponential. The number of hidden units is varied to parameter match the models.

**Figure 6:**
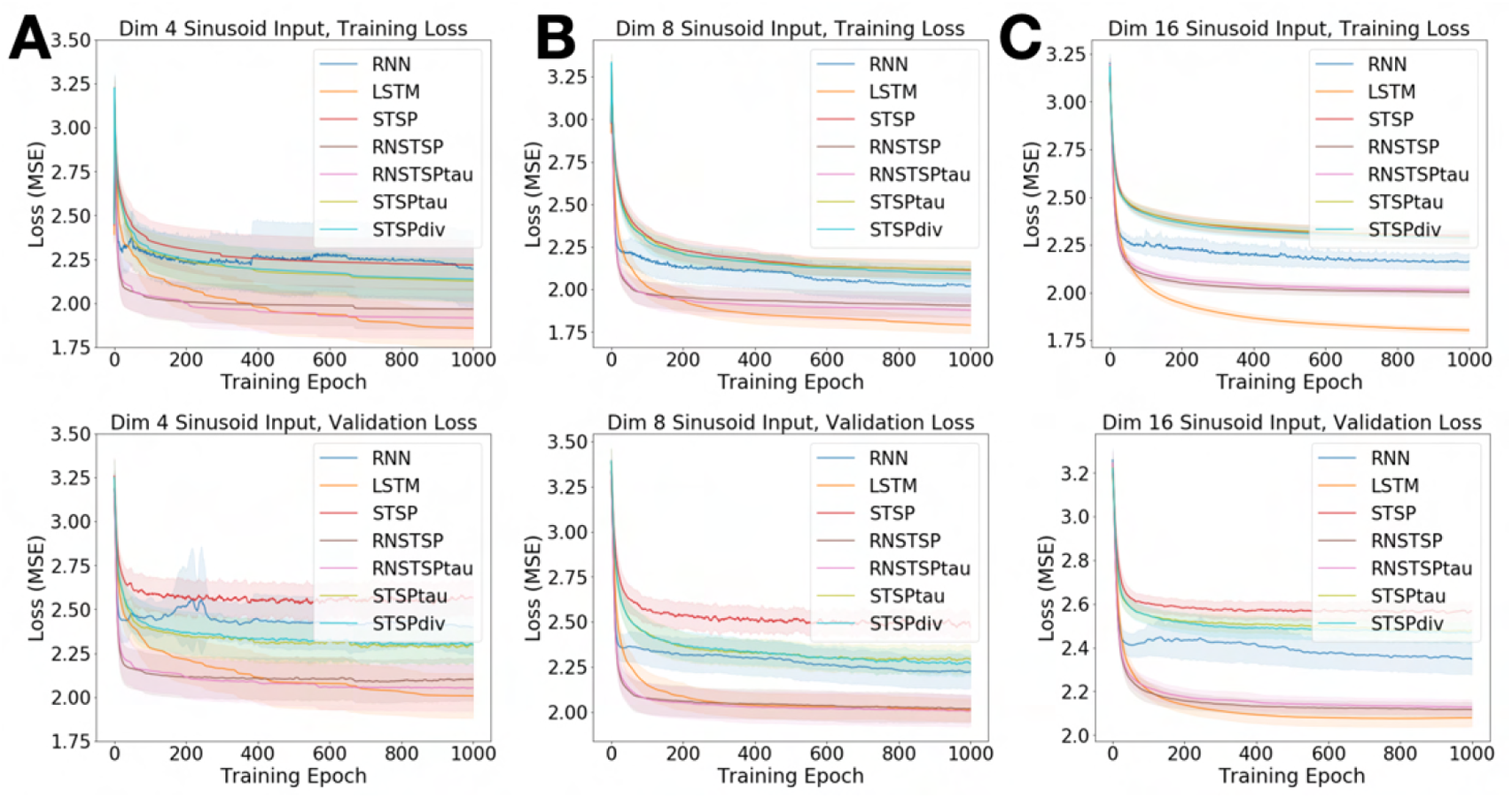
Training (Top) and validation (Bottom) loss across training epoch for models predicting a 4 **A**, 8 **B**, and 16 **C** dimensional sinusoid when the number of hidden units is fixed at 16 nodes. Under these conditions the LSTM model has significantly more trainable parameters than the other models (Table 4).

**Figure 7:**
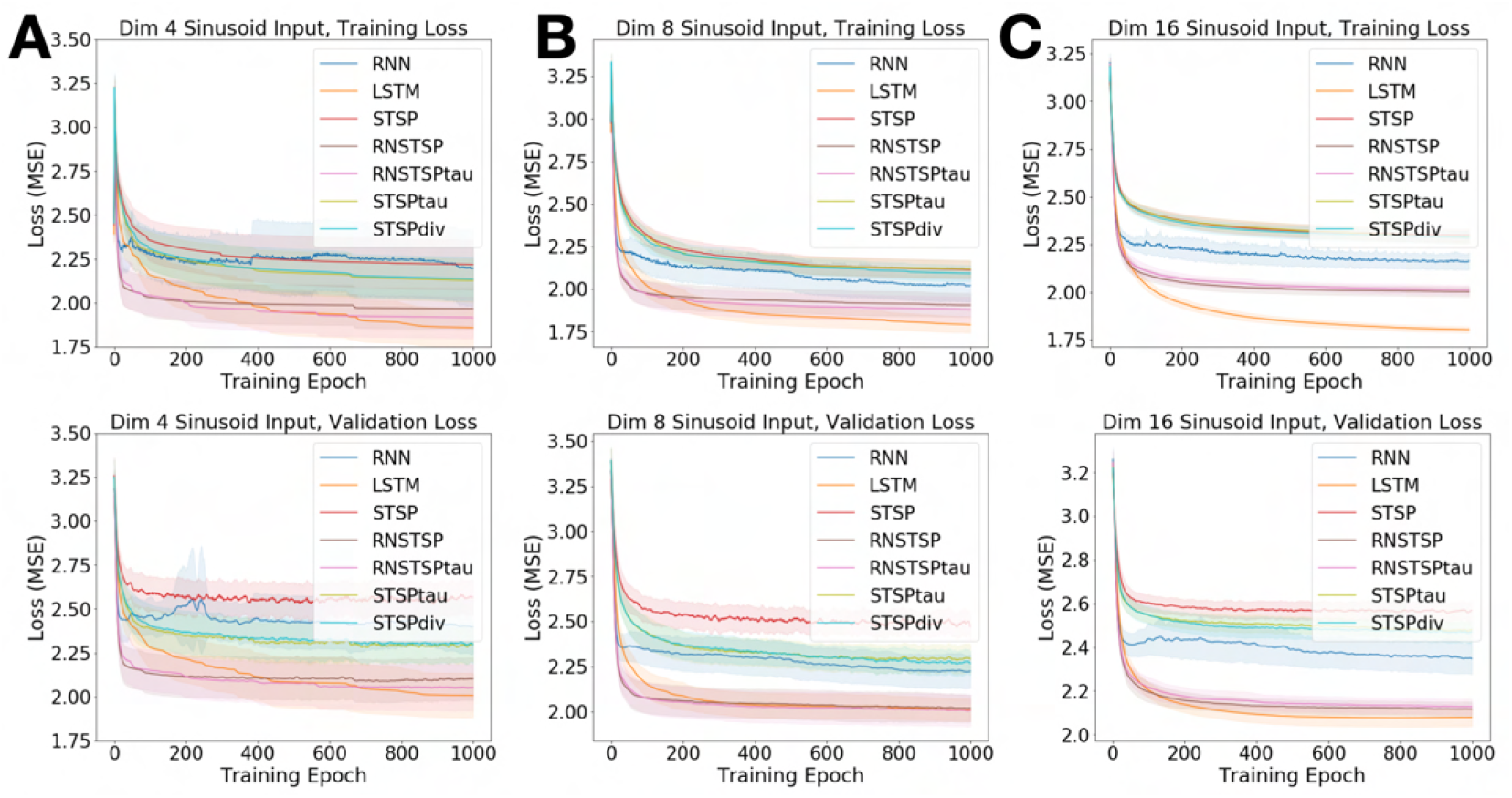
Training (Top) and validation (Bottom) loss across training epoch for models predicting a 4 **A**, 8 **B**, and 16 **C** dimensional sinusoid. The sinusoidal dynamic learned in this case differs from that of Figure 1 and Figure 4 in that the frequencies of oscillation are instead randomly selected from a uniform distribution from 0.001 to 0.333 Hz. The number of hidden units is varied to parameter match the models.

**Figure 8:**
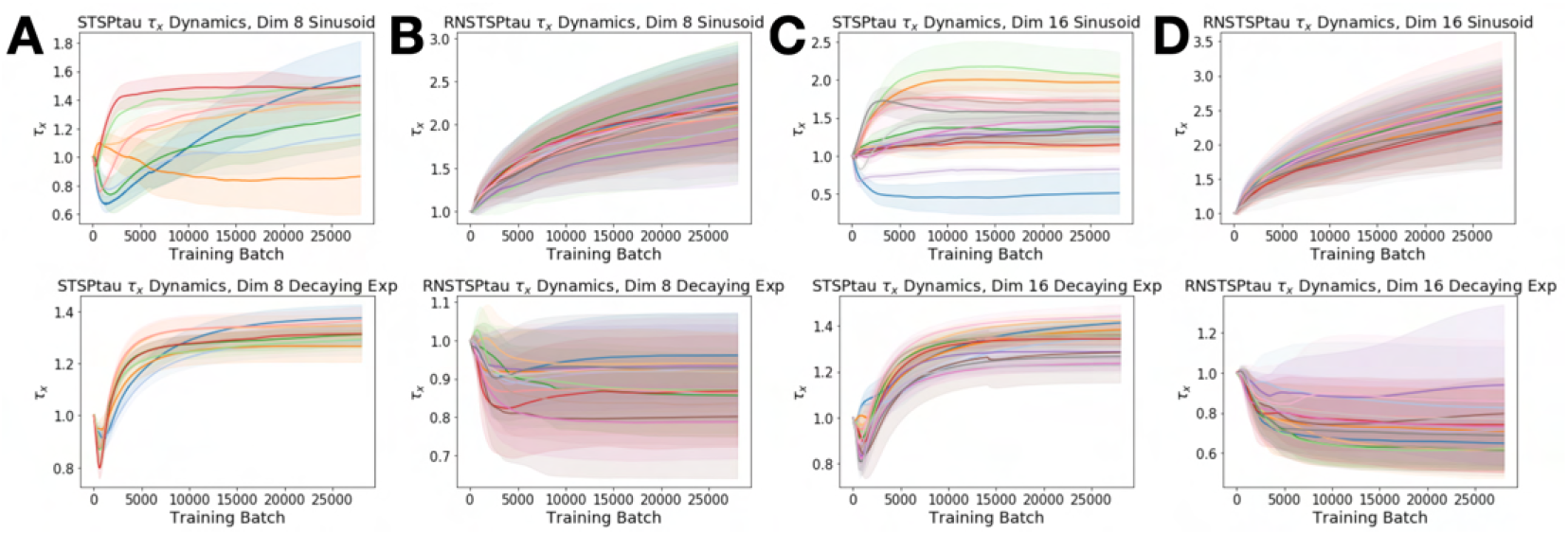
Charts demonstrating the heterogeneity of synaptic dynamic time constants for the RNSTSPtau and STSPtau parameter matched models trained on 8 and 16 dimension sinusoid (Top) and decaying exponential (Bottom) inputs. The number of STSP nodes differs between the STSPtau and the RNSTSPtau models because the nodes with STSP are in the input layer for the STSPtau model versus the hidden layer in the RNSTSPtau model. **A** STSPtau time constant evolution across training for dimension 8 sinusoid and decaying exponential inputs. **B** RNSTSPtau time constant evolution across training for dimension 8 sinusoid and decaying exponential inputs. **C** STSPtau time constant evolution across training for dimension 16 sinusoid and decaying exponential inputs. **D** RNSTSPtau time constant evolution across training for dimension 16 sinusoid and decaying exponential inputs.

**Figure 9:**
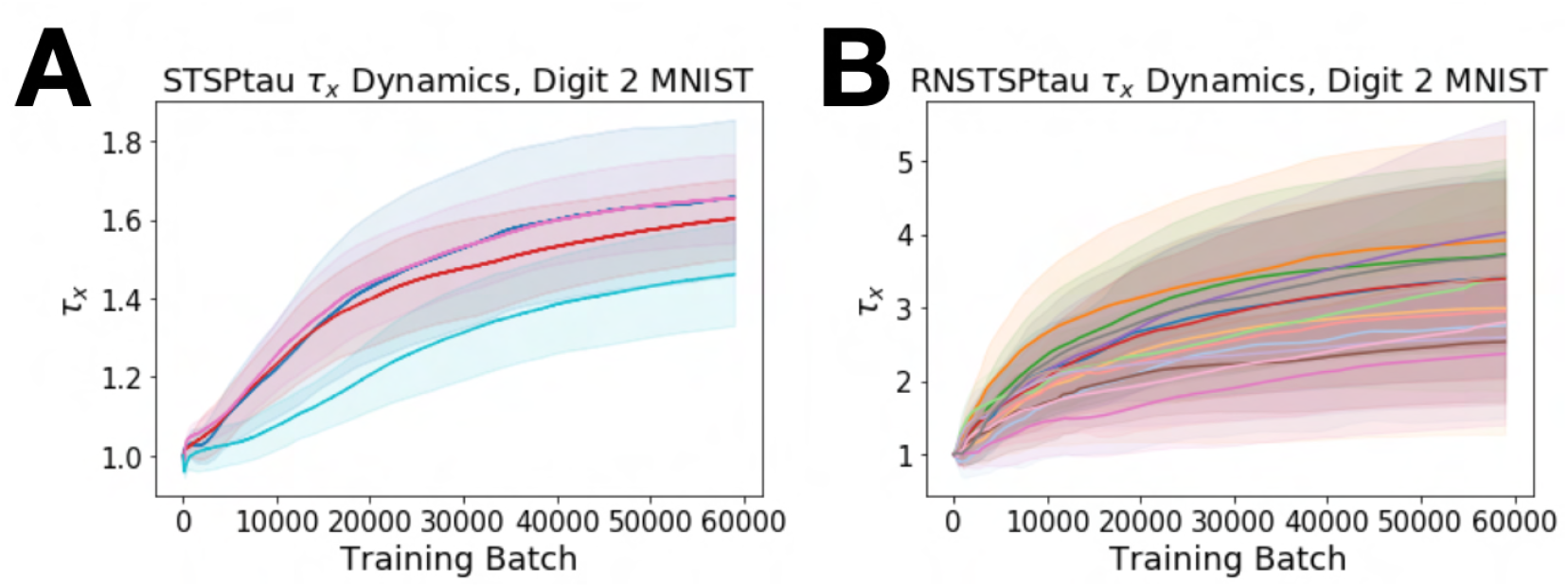
Evolution of synaptic dynamic time constants across training for the **A** STSPtau and **B** RNSTSPtau parameter matched models trained on the sequential MNIST task.

**Figure 10:**
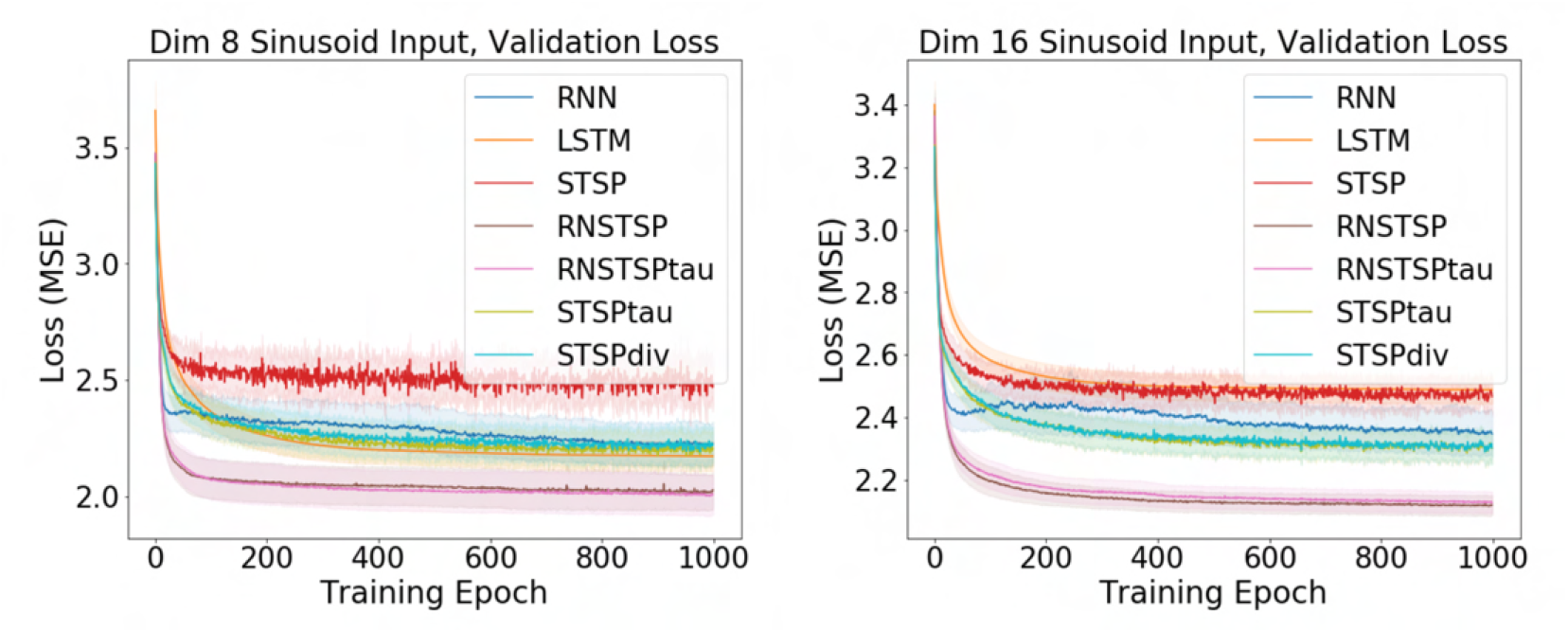
Validation loss plots for the dimension 8 and 16 sinusoidal prediction tasks that are identical to those presented in Figure 1C, D but without moving average smoothing.

## Notes

### Competing Interest Statement

The authors have declared no competing interest.

